# Morphological integration between seeds and seedlings of *Hymenaea courbaril*

**DOI:** 10.1101/2020.04.10.036178

**Authors:** Leonardo de Lima Pereira Regnier

## Abstract

Morphological integrations are unknown to forestry seeds. Understanding if seed measurements could predict its future seedlings features is a central question in seedling production. *Hymenaea courbaril* is an important species in this context and to the applied forestry. Thus, this study aimed to understand how some seedling features, could be related to the seed weight, and be affected by the population origin. The measurements consisted of seedling collar diameter, weight, protophilus area, central and lateral vein. Seed weight consistently varied between the populations in the study. Both populations had higher weight ranges than mentioned in the literature. There was no strong evidence that greater seed weight requires lesser time to germinate, conflicting with previous information. All the measurements presented enough shreds of evidence to be considered different when comparing the populations, except for the protophilus area and lateral vein length. All the studied measurements presented low correlation indexes to seed weight, except for seedling collar diameter, and seedling weight, which presented a moderate correlation. Protophilus elongation pattern was strongly associated with the leaf width when compared to midvein.

## INTRODUCTION

Bigger seeds have greater food supplies, which could result in greater initial growth rates [1]-[4]. In order to identify the best propagation methods, seed size and weight have been used to identify seed lot quality in several plant species [1], [4], [5]. Some studies have shown that biometrics of seeds could be used as an indirect parameter to propitiate grater and homogenous germination, and also favoring the germination of the embryo resulting in the most vigorous seedlings [1], [4], [6]. In general, seeds displaying larger size or density are potentially more vigorous because they usually have more reserves and they present well-formed embryos [7].

Seed mass is a feature of great importance because it is linked to recruitment via germination, seedling establishment, and offspring, [7]. There is some uncertainty about the relationship between seed weight and/or size in seedling vigor [7]. While crop science has been shown that, to economically relevant species, seed size is an appropriate parameter to measure seedling quality and vigor, in forestry species case, and especially from tropical countries [4], this is not extensively explored [8], [9].

Studies focusing on the morphometric relationships of bushlands species are extremely relevant to recognize them in the field, and also to understand if these parameters reflect better seedling quality, which would enable us to employ this knowledge in recovery programs and urban afforestation [4], [10].

*Hymenaea courbaril* L., popularly known as *“Jatobá”* is a deciduous tree [11], with a wide geographic distribution, from southern Mexico to southeast Brazil [12]. This species has a high tolerance to great climatic and hydric variations and also low nutritional requirements [13], [14]. Moreover, this species belongs to the climax ecologic role [15], with rare occurrence (less than 1 (tree/hm^2^), and sparse distribution [14].

*H. courbaril* seedlings are heterophilic, with protophilus anatomy divergins from the other leaves. The protophilus of this species is characterized by the presence of stipules, rounded apex, and an expressive primary veining, with a full edge, and ovoid format [16].

Compared to other biologic taxon, plants are the least explored in morphological modularity, and developmental approaches in the search of modules are even rarer [17], [18]. Notwithstanding, because of their modular body plans, the plants offer additional opportunities to study integration among structures within and among individuals or populations [19].

Morphological characterization and influence on seed and fruit quality have been scarcely explored to forestry seeds [9]. In this context, *H. courbaril* has the potential for quick production and high seedling survival in the field [11], [20]. These features favor faster seedling production and establishment in the field, which also promotes the regular use of this species in forestry and afforestation programs [20]. Besides the relevance of studies concerning biometric relations and influence on germination of this particular species, they are extremely rare [1].

Therefore, this study aimed to understand how seed weight could be related to the main seedling features of two populations of *H. courbaril.* In this study particular case, we evaluated the collar diameter, weight, protophilus main dimensions, and area.

## MATERIAL & METHODS

This study was conducted in the city of Cotia (23°36’30.0”S 46°50’48.9”W) between the years 2019 and 2020. The region presents the Cwa climate, altitude tropical climate. Characterized by dry winter and concentered rains during summer, according to the Köppen climate classification [21]. Described mainly by dry winter, and the highest mean temperature above 22°C.

Plant material was harvested on 31 October 2018, from two populations, 20 km away from each other, used to measure variations associated with seed origin. Only mature fruits from the ground were gathered. The seeds were obtained by shattering the fruits with a hammer. The endocarp was manually grated with a blunt knife.

Each seed had its weight measured and received an identification number to enable tracking of the formed seedling (Figure 1). Seeds were planted in sterile thin quartz sand (1 mm particles) and kept in colorless polypropylene boxes. During the analysis period, each box received a pulverized 500 mL water supply twice a day at 8 am and 5 pm. Emergence was recorded for 75 days. All seedling measurements (collar, first leaf pair area, seedling weight) were performed with fresh seedlings at the 21° day after the radicle protrusion, in order to minimize the effects of latter germinations [6]. At the end of the study period, the sample consisted of 380 seedlings from population I and 255 seedlings from population II, 635 individuals in total.

**Figure 1:**
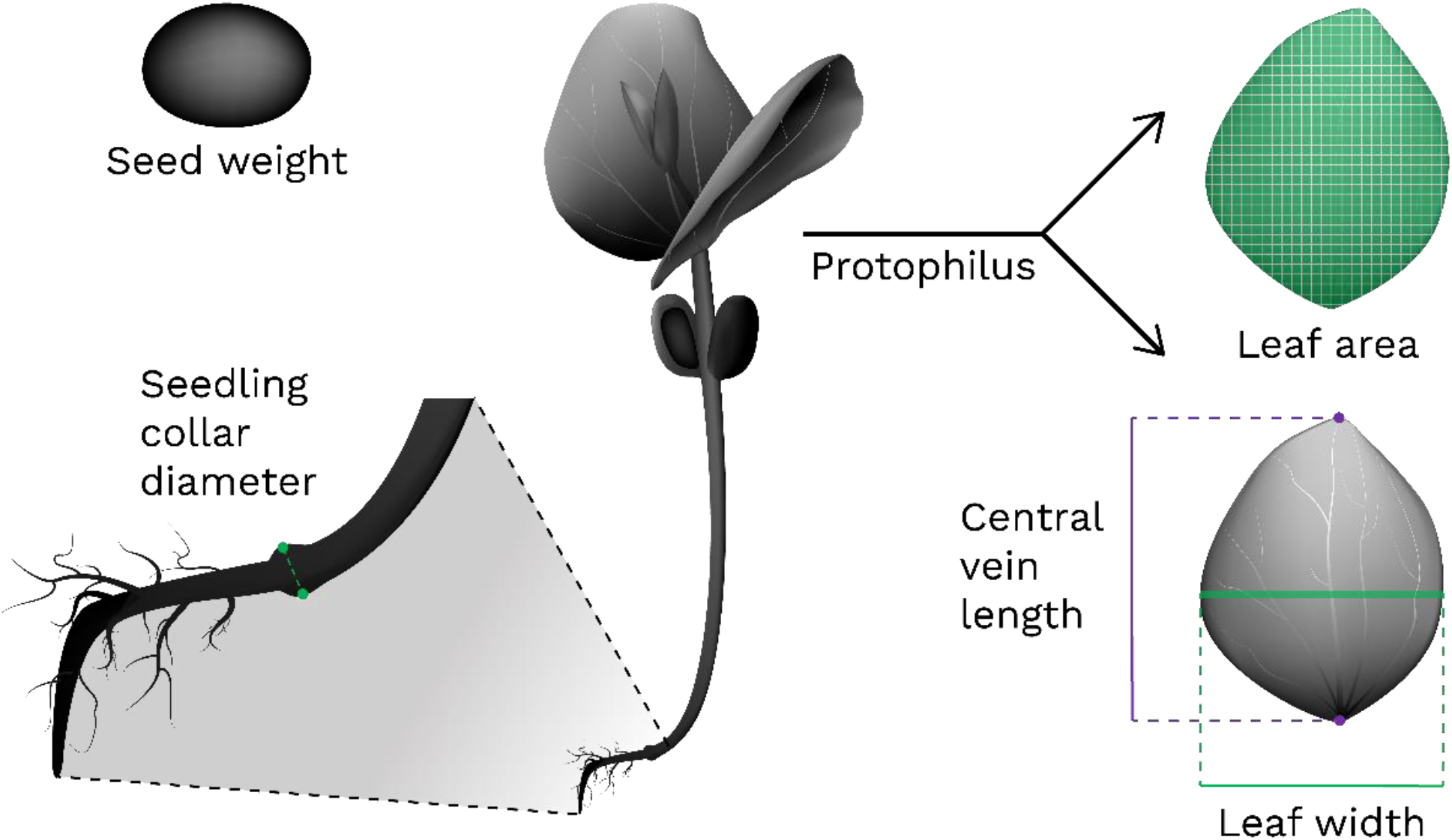
Studied seed and seedling measurements. After the establishment of seed identity, the seed weight, and it generated seedlings had the collar diameter, protophilus area, and this leaf linear dimensions: central vein (midvein or primary vein) and lateral vein (leaf width) measured [21].

Seedling vigor was graded through the measurement of the first leaf pair (protophilus) area, principal vein, collar diameter, and seedling weight (Figure 1). One of the protophilus was glued to a paper sheet and scanned. The protophilus area was measured using Paint.net^®^ software [22], using the “magic wand” tool, and the principal vein length and width were obtained through the “line/curve” tool, drawing over the midvein and the largest width dimention of the scanned leaf.

The collar diameter was measured on the mild dilatation of the neck [16] with a digital caliper. Seed and seedling fresh weights were measured using a digital scale. The seedling fresh weight was always performed after morning irrigation at 8 am.

Boxplot associated with violin plots were obtained through R software [23] with vioplot [24], and ggplot2 [25], using the dplyr [26], and magrittr [27] packages to deal with the amount of data. The density plot was constructed also using the ggplot2 [25] package.

Correlation graphics were obtained from the Excel component of the Microsoft Corporation Office pack and represented through scatter plots, and regression lines were chosen according to the correlation index. The model with the best fit was chosen to be displayed. Principal Component Analysis (PCA) was performed with the R software [23], first with FactomineR [28] and in sequence, the biplot was performed with factoextra [29] package. All statistical tests were also executed with R software [23] using agricolae [30] and GerminaR package [31]. The results were tested by ANOVA and in sequence submitted to Tukey test with a critical p-value of 5% (p <0.05).

## RESULTS AND DISCUSSION

### Seed weight and its generated seedlings

The experimental seed weight distribution showed differences in the general pattern between both populations (Figure 1). Population II presented greater amplitude and variable frequencies of seed weight. Having many seeds between 3 and 5 g and other high frequencies between 5.5 and 6 g, while Population I had almost the usual normal distribution, with the most frequent value about 6.5g, and with more concentered values of seed weight.

In the present study, we obtained a minimum of 2.7 and a maximum of 8.7 g in both populations (Figure 2). Based on the seed weight classification of other studies with *H. courbaril*, these seeds would be considered big [8]. Cruz & Pereira [32] found a mean seed weight of approximately 2.85 g, And some authors point to a weight range of 2.1 – 6.5 g per seed [15]. Other authors have indicated that the most common mean weight of the seeds of *H. courbaril* is approximately 3.4 g [16], [33]. Silva et al. [34] presented a slightly wider range from 2.62 to 4.36 g. But consistent wider ranges could be found in Andrade et al. [35], which found seeds from 4.08 to 11.2g. Those last authors also mention that moisture content could also be responsible for the variations between seed mass, which was still indicated in other previous studies [36]. The storage time [12] and the harvesting method [36] were also indicated as factors that could affect the seed mass.

**Figure 2:**
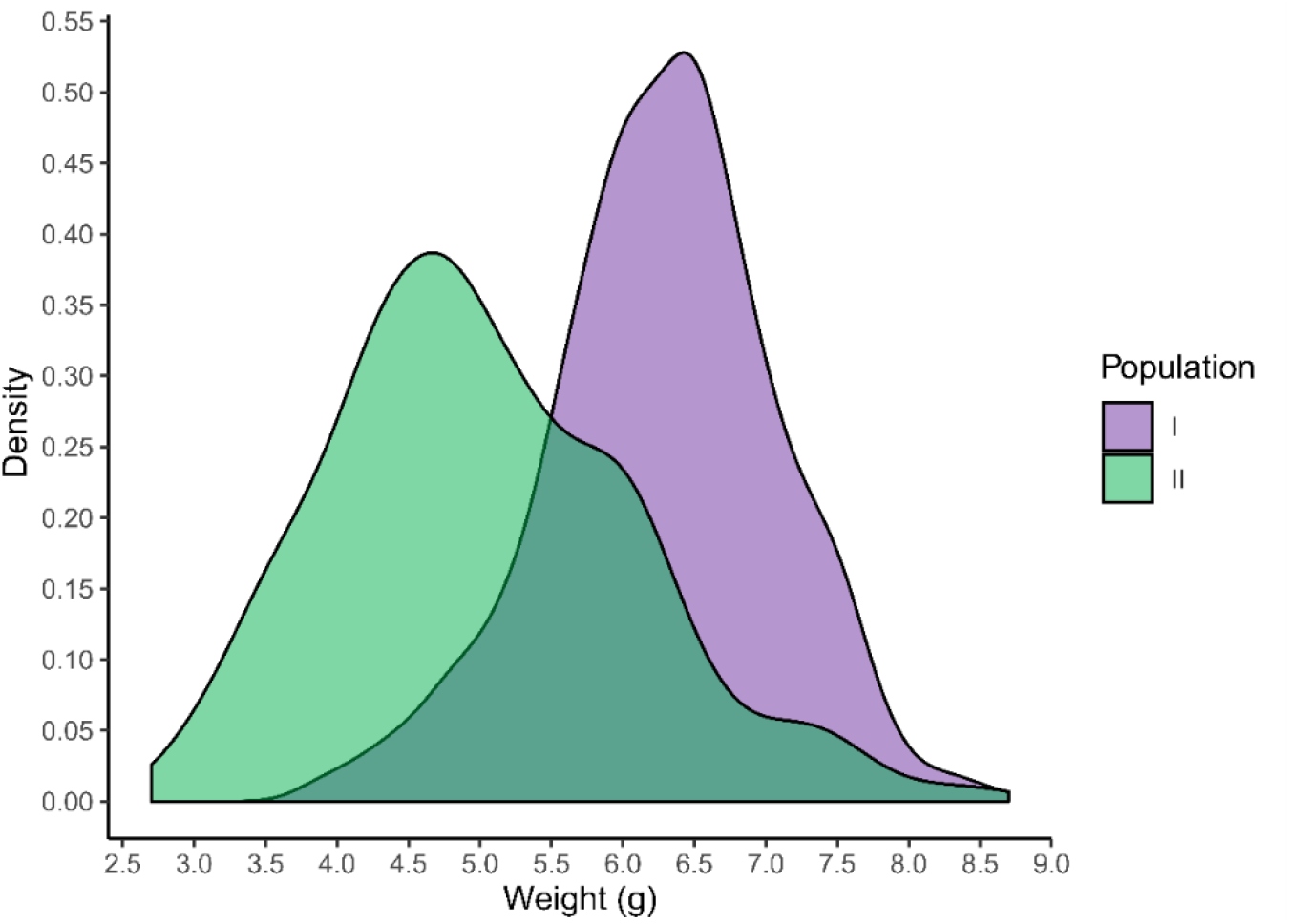
Density plot of *Hymenaea courbaril* seed weight distribution according to origin: Population I 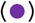 and Population II 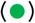.

Tropical arboreous species usually present a high variability in seed mass [4]. Other studies mention that seed weight is highly variable due to the water content [35], which also varies according to some environmental conditions during seed development, and features, such as maturation stage [36], [37]. However, this species seed moisture content could vary between 12.46 until 16.05 % of the weight [37], a very strict range in view of the diversity of values found to seed weigh disposable in the literature, or even the present study variation. In addition.

Several factors are pointed out as responsible for this sort of variation. Some authors have indicate that seed mass is controlled predominantly by genetic information from maternal and zygotic tissues. Thus, the great genetic diversity of these species [38], [39] is generally pointed out as the mean cause. However, seed growth is also influenced by environmental cues [7], [9], [40] such as edaphoclimatic differences [9] and ecologic interactions [4].

Statistically analyzing seed weight, we found remarkable differences between the populations (Figure 3). Most seeds from population I have seed weight concentered in 6.275 g, a 25% greater than the population II mean.

**Figure 3:**
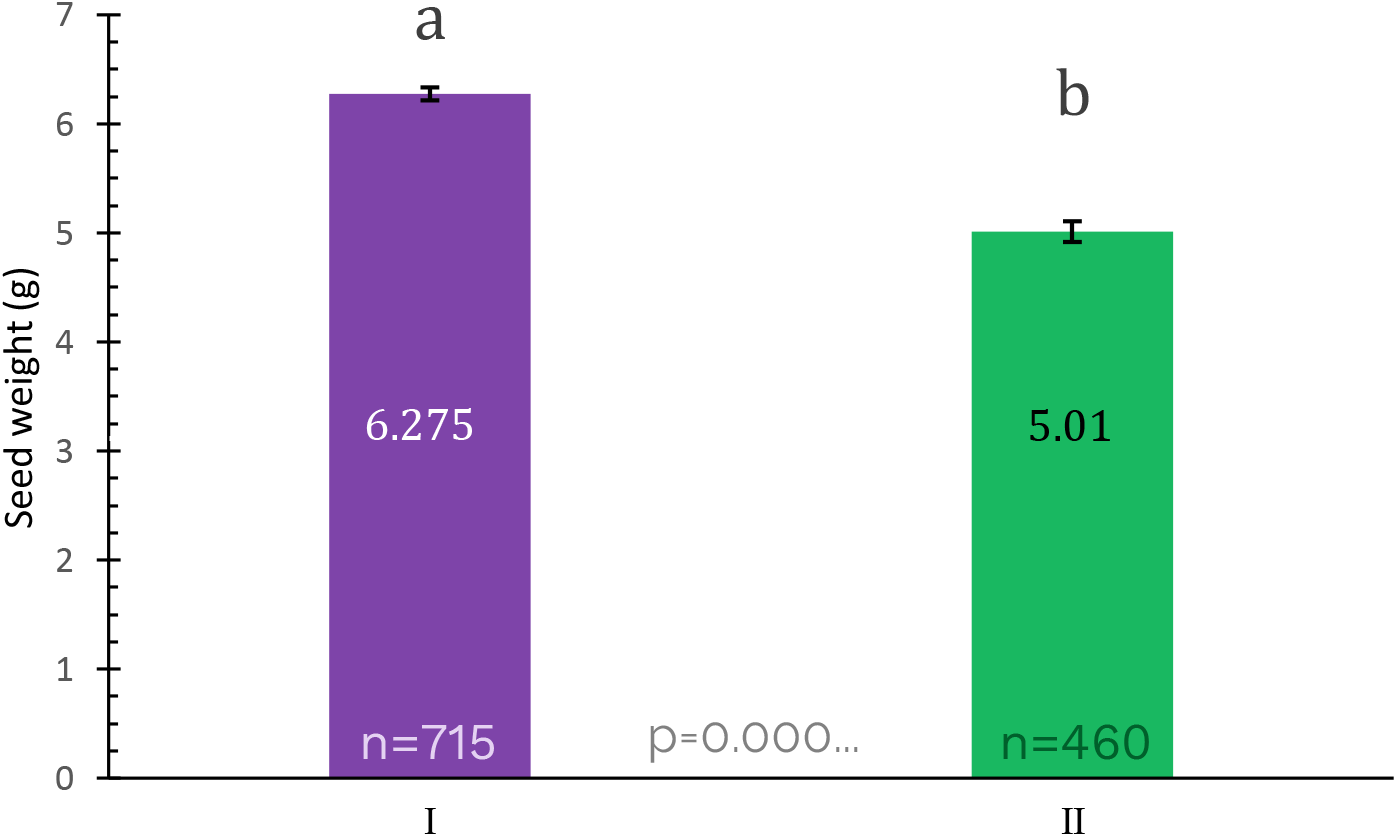
Mean seed weight of *Hymenaea courbaril* according to origin: Population I 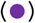 and Population II 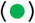. Error bars present a confidence interval of 95%. Different lowercase letters present statistical differences at p<0.05.

Even if we try to use other central measures, such as median or mode, the obtained values are still very close to the mean. Obtaining, to these indexes, variations of 0.2 g from the mean values. The same result was observed for population II, keeping the value close to 5.01 g. This great consistency of the central indexes is obtained due to the ample sample used in this study. Corroborating with previous studies conducted with these populations [36], which already presented similar mean values of seed weight.

In the seedling production context, seed weight is usually a parameter to measure the seed lot quality and genetic diversity [1], [4], [9]. In addition, it is relevant, as these seeds are usually sold per kilogram [9], [41], [42].

There is still conflicting information concerning the influence of origin on germination and seed parameters. Variations between seed weight of different parent plants have already been mentioned in the literature [9], [32]. Other authors concluded that even when these differences were present, they did not provide substantial variations in germination aspects [39], [43].

It is not completely understood which factors could lead to variations in seed weight. Although it is already recognized that seed weight is related to parent plant height, especially in Fabaceae [44]. Similar evaluations of seed dimensions, such as weight, were conducted and seeds from different parent populations had a significant morphometric difference, but on the other hand, the seed weight between them was almost the same [9].

### Seed weight and germination

Correlations between seed weight and the time required for germination is a very usual analysis [4]. The results indicate that the correlation to seed weight and time to germinate has a weak correlation value (R^2^=0.1) [45], in population I, II, and even when considered both of them (Figure 4). The correlation lines also differed between the populations, while population I had an almost steady value, with a low correlation index, population II had a positive correlation and substantially higher coefficient of determination (Figure 4).

**Figure 4:**
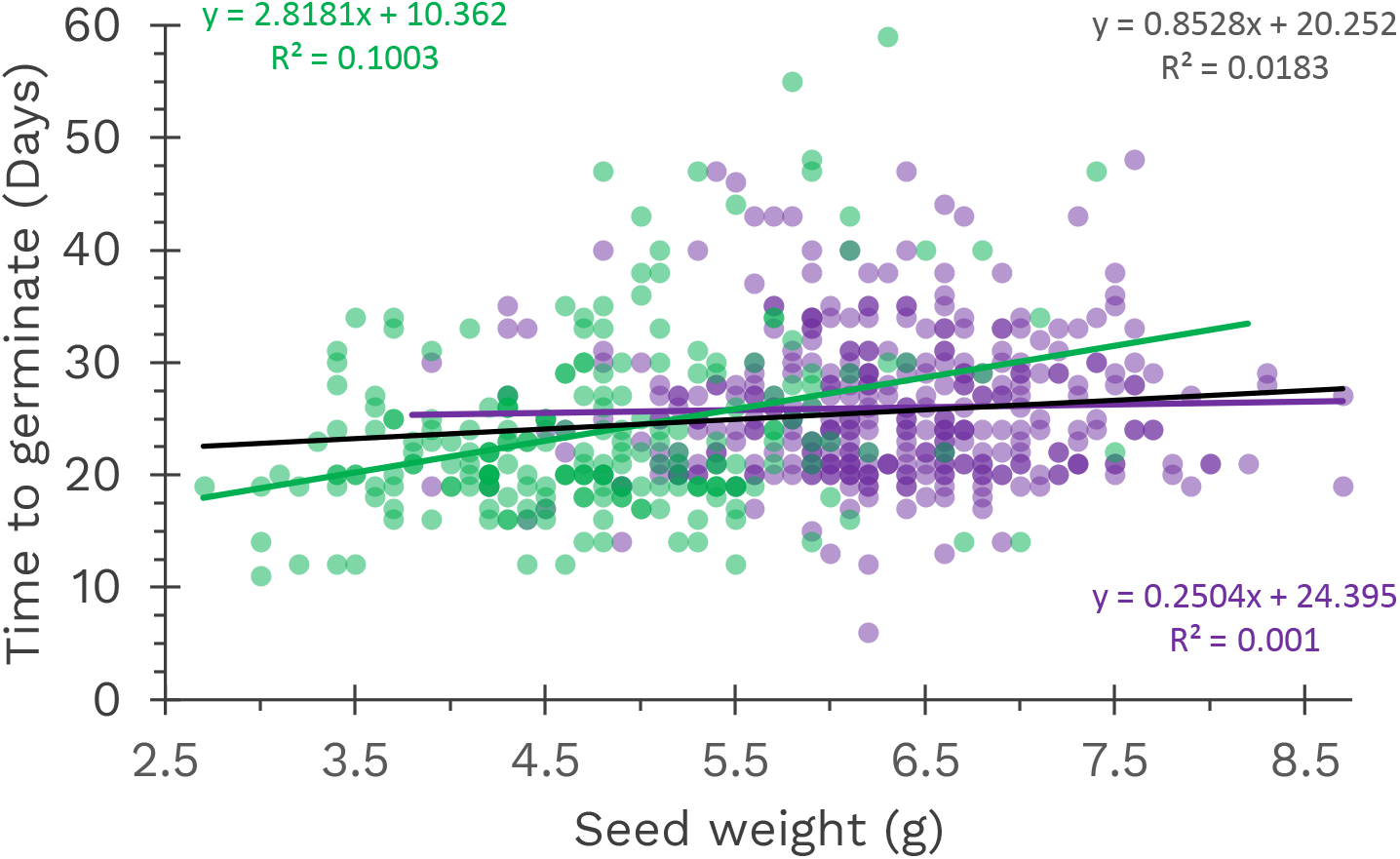
Scatter plot with regression lines of seed weight and the required time to germination of *Hymenaea courbaril* according to origin: Population I 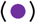, Population II 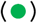, and both 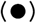.

Previous analysis indicated that greater seed weight promoted faster germination rates to *Parapiptadenia rigida,* another member of the Fabaceae family [7]. When considering the populations isolated, to population I, the relationship between seed weight and time required for germination is positive. Indicating that the seeds of greater weight required greater time to germinate (Figure 4). This result is the opposite of that mentioned for *P. rigida.* This result also emphasizes that seed vigor and responses are best examined using the population perspective [6].

Both populations presented a weak association between the variables (Figure 4). Other studies, with another species from Fabaceae, also did not find associations between seed weight and seedling vigor [5], [46]. Seedling survivor and faster germinations associated with seed weight were also not observed [4].

Considerable discrepancies in germination proportion, germination speed, and mean germination time between populations to *H. courbaril* were found by some authors [9]. Endosperm growth is pointed as the main cause, because it could be affected by parental origin [47].

However, recent studies have not found consistent variations between the germination indices associated with different populations [43]. Thus, it is still not clear what truly affects seed weight variations between the populations, requiring substantial analysis in the future.

### Seedlings measurements

The most common parameters to evaluate seedling vigor indicate that population origin substantially affected most of the studied measurements, except for the lateral vein and protophilus area (Figure 5). For some variables, there were great dissimilarities between the studied populations. To seedling collar diameter, for example, there is almost a bimodal distribution to population II (Figure 5-D).

**Figure 5:**
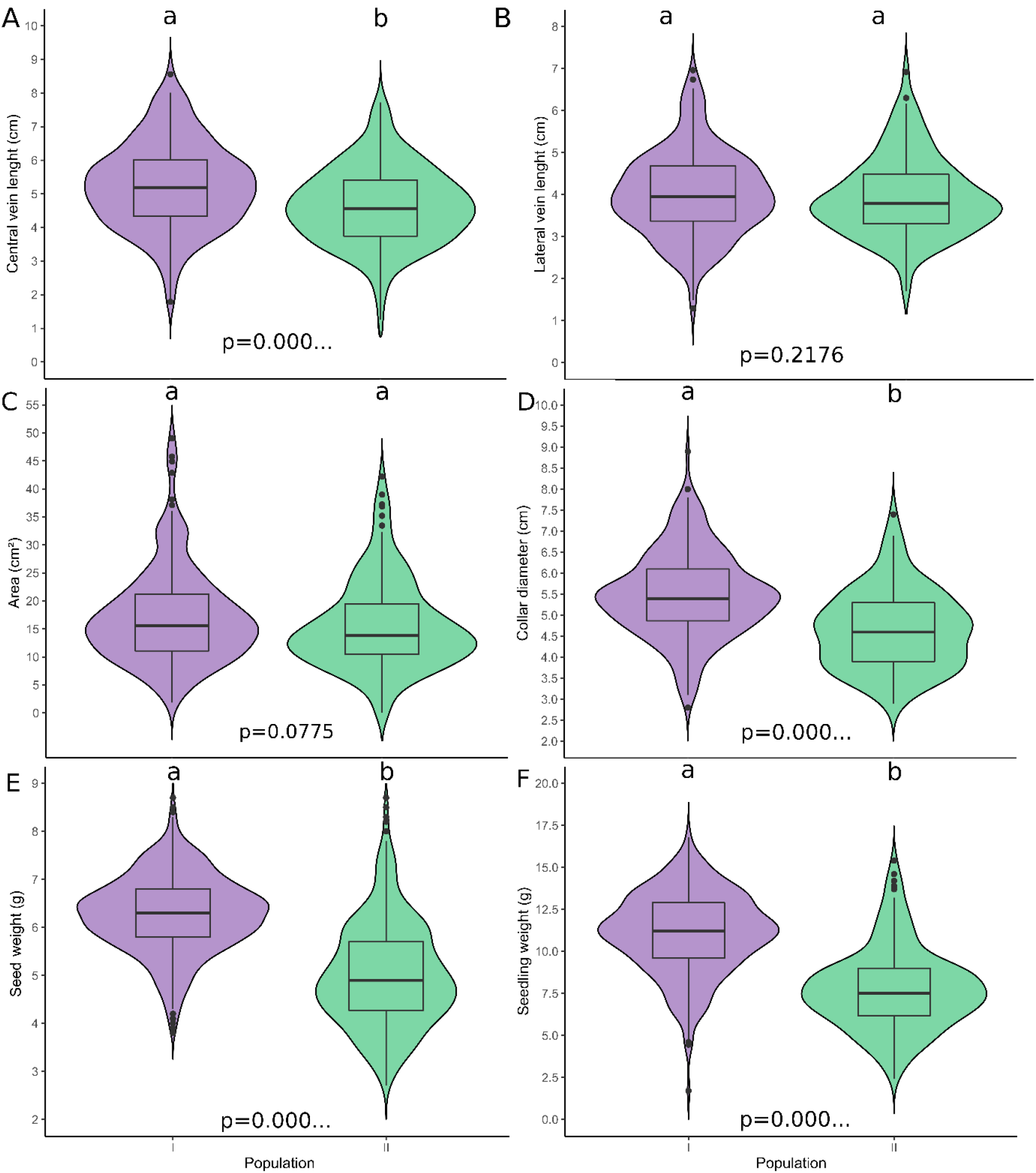
Box and violin plots of seed and seedling of studied measurements of *Hymenaea courbaril:.* **A** – Protophilus central vein (midvein); **B** – Protophilus width; **C** – Protophilus area; **D** – Seedling collar diameter; **E**- Seed weight, and **F**- Seedling weight. According to origin: Population I 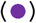 and Population II 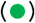. Lowercase letters present statistical differences using α≤5%.

The analyzed measurements are important in the seedling production context because it is necessary to generate fast and uniform germination and also homogeneous seedling size [47]. Usually, in the literature, the fact that populations with lower values of seed weight generated smaller seedlings will be erroneously associated as a causal association, since there was no proper analysis to establish this relation. Although, these feature correlations will be better discussed in the section **Seedling morphological integrations**.

Protophilus, as the first photosynthetic leaf pair of this species, has important implications related to its dimensions and area because it is generically associated with the survival capacity of the embryo. Given that, before protophilus development, the seedling is dependent on the cotyledon reserves [16]. The data indicate that consistent divergences were only found in the protophilus central vein, while to the lateral vein and area there was no difference. This is possibly related to the strict relation between the protophilus area and width length. Considerations of leaf elongation patterns are explored in the last section of this paper.

Ferraz & Engel [48], found that collar diameter during development could range from 3.94 to 4.37 mm. The mentioned range is consistent with the values found for population II but very distant from the values of population I (Figure 5-D). In the literature, it is also possible to find the opposite scenario, in which the given information is consistent with population I but not to population II when the study data is compared to the mean values we found. For instance, the mentioned by Costa et al. [20], in which the seedling collar diameter raged from 4.9 to 5.07 mm.

The seeds and seedlings from population I were consistently lighter and smaller in almost all the studied measurements, except for the lateral vein, and the protophilus area (Figure 5), probably due to the association of these variables.

It is important to compare populations to comprehend the multiplicity and infer the remaining genetic diversity between them [9]. In general, variations between populations in seed and fruit morphometrics are expected because they are under different environmental conditions [9], [42], and evolutionary processes [38], [42]. Part of phenotypical measurements could also be due to plastic responses to heterogeneity in environmental factors [19].

Thus, it seems that the studied populations still present some level of individual variability, presenting discrepancies to some of the studied dimensions. Besides that, in germination aspects, these populations did not present significant discrepancies [39].

### Seedling morphological integrations

The possible correlations between seed weight and the protophilus studied dimensions, area, and seedling collar were very low (Figure 6). The best correation could be obtained between the seed mass and seedling collar diameter (Figure 6-D), but even in that case, which had a potential pattern of a positive correlation of the variables, only 7.66% of the obtained variation could be explained by the regression line.

**Figure 6:**
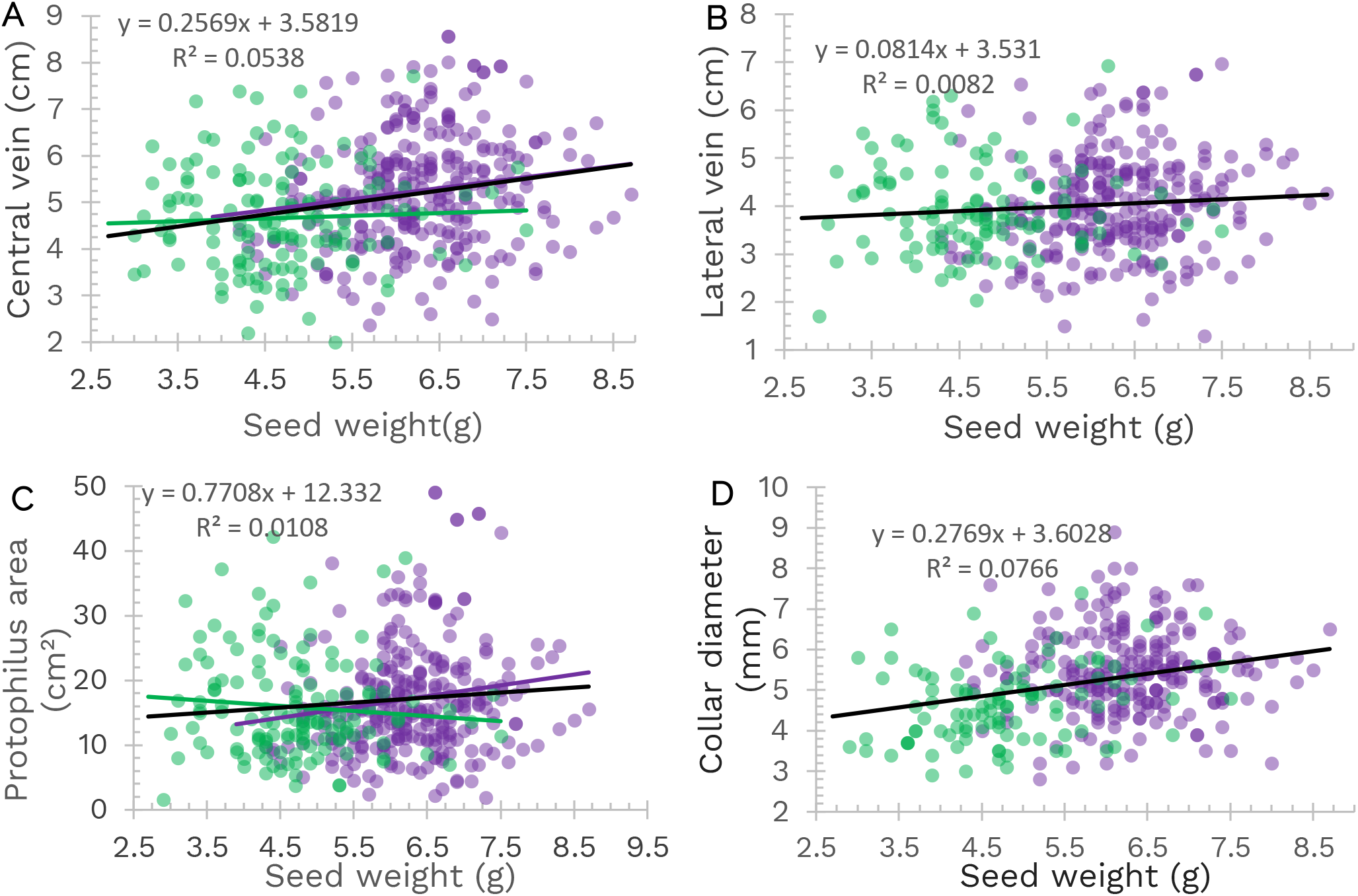
Scatter plot of seed weight and seedling measurements of *Hymenaea courbaril*: **A** – Central, vein, **B**- Lateral vein, **C**- Protophilus area, and **D**- Seedling collar diameter. Colors represent the seed origin: Population I 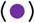, and Population II 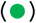. General regression lines and their coefficient of determination (R^2^) are also presented.

In general, both populations had the same tendency, in that case, only the regression line using the data obtained by both populations was presented (Figure 6-B and D). Although, for central vein dispersion, population I indicated a positive correlation to seed weight, while population II presented almost no relation, with a steady tendency (Figure 6-A). Similar patterns were observed in the protophilus area, but in that case, the populations presented an opposite association, population I indicated a positive, and population II a negative correlation (Figure 6-C). The regression line of both data had a slight positive association between seed weight and protophilus area (Figure 6-C).

It is expected that large seeds could promote greater leaf area production, which was already observed in some species, especially at the beginning of the growth cycle [49]. In the present study, the results to *H. courbaril* do not corroborate this hypothesis, since higher seed weight does not correspond to greater protophilus area, or any of the studied protophilus dimensions (Figure 6-A, B, and C).

Also to this species, Pagliarini et al. [47] mentioned that higher seedlings are not related to larger stem diameter. The present data also emphasizes that there is no clear relation between seed weight and seedling collar diameter. Other studies have shown that the environment could also affect morphometric evaluation. Hydric stress seems to affect the seedling leaf area [13], and light exposure could also influence some seedling measurements, such as the collar diameter [20]. Hence, it is possible that studies using different conditions could also find variations in the results mentioned here.

In the morphometric analysis, an important facet is the continuum relations between the variables and not only classification as all-or-nothing [18]. These study results indicate that the seedling collar diameter, the protophilus dimensions, and the area is not strongly linked to seed mass. Thus, to *H. courbaril*, it is not useful to estimate seedling vigor, or the studied dimensions, based only on seed weight.

Seed and seedling weight had a moderate positive association (Figure 7) [45]. Seeming that there is, somehow, a relation between these variables, in which seeds with greater weight had greater seedling fresh weight (Figure 7).

**Figure 7:**
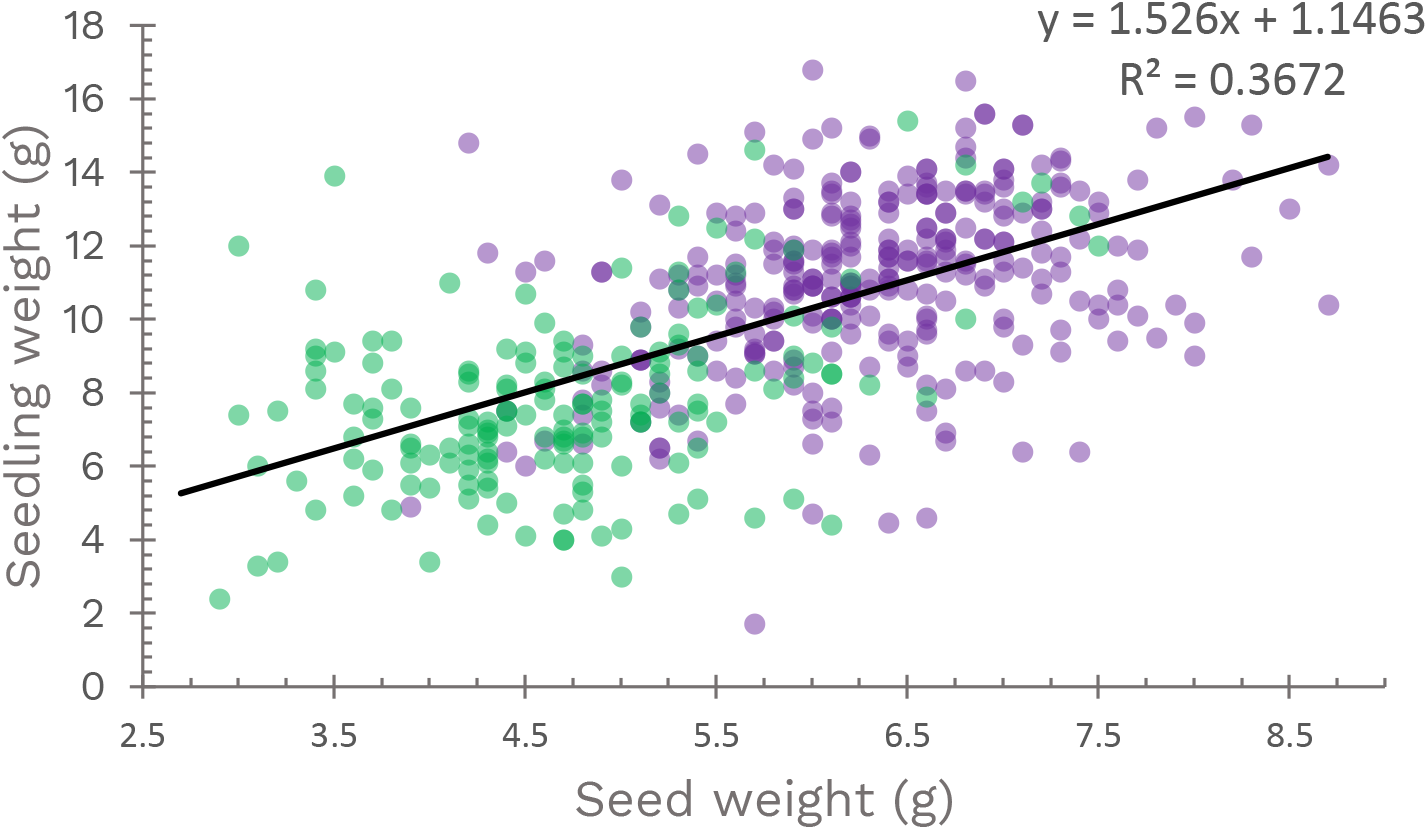
Scatter plot of seed weight and seedling weight of *Hymenaea courbaril.* Colors represent the seed origin: Population I 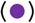, and Population II 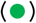. The general regression line and its correlation indexes (R^2^) are also presented.

Recently, knowledge of the morphometric relation between seeds and seedling vigor has been raising interest, especially to identify features of the seed that reflect its quality, or more vigorous seedlings. [4].

In usual production practices, poor seed vigor can have direct and serious financial implications. Because low-quality seeds require labor, glasshouse space and/or planting materials, and results in a low-quality product, or non-uniform production [6]. These motives make understanding morphological integrations crucial to seedling production.

Identifying the influence of seed weight on seedling vigor seems to be specific to some species and restricted to some groups [4]. In general, studies that try to evaluate those relations use a discrete approach to classify seeds as “small”, “medium” or “big”, this sort of classification besides subjective, and restricted to the sample amplitude, reduces the data resolution [45].

It is somehow expected that seed and seedling weight had some kind of relation because on the 21st day of study, the cotyledons are still attached to the seedling, as described by Duarte et al. [16].

Development associations seem to be less determinant than associations through organ function in plants, but a proper investigation is still needed [17]. Parent plant height seems to be correlated with seed weight in some families, Fabaceae is one of them [44]. Previous studies found that the group with bigger seeds presented higher germination rates I8]. At the same time, differences in germination speed and proportion to *H. courbaril* were not found in parent plants 30 meters [43], or populations 20 kilometers away from each other [39]. These discrepancies could result from the chance since the sample of some studies was very small, the development stage used in the analysis is not always the same, and also the approach of dealing with continuous variables as they were discrete [45]. Furthermore, direct interaction between developmental pathways that are sensitive to the same outside stimulus could also respond, resulting in covariations, besides the lack of integration [18], [19].

The measurements associated with the protophilus, especially the central vein and area, were the most variable feature between the individuals (Figure 8). Being responsible for most of the data variation.

**Figure 8:**
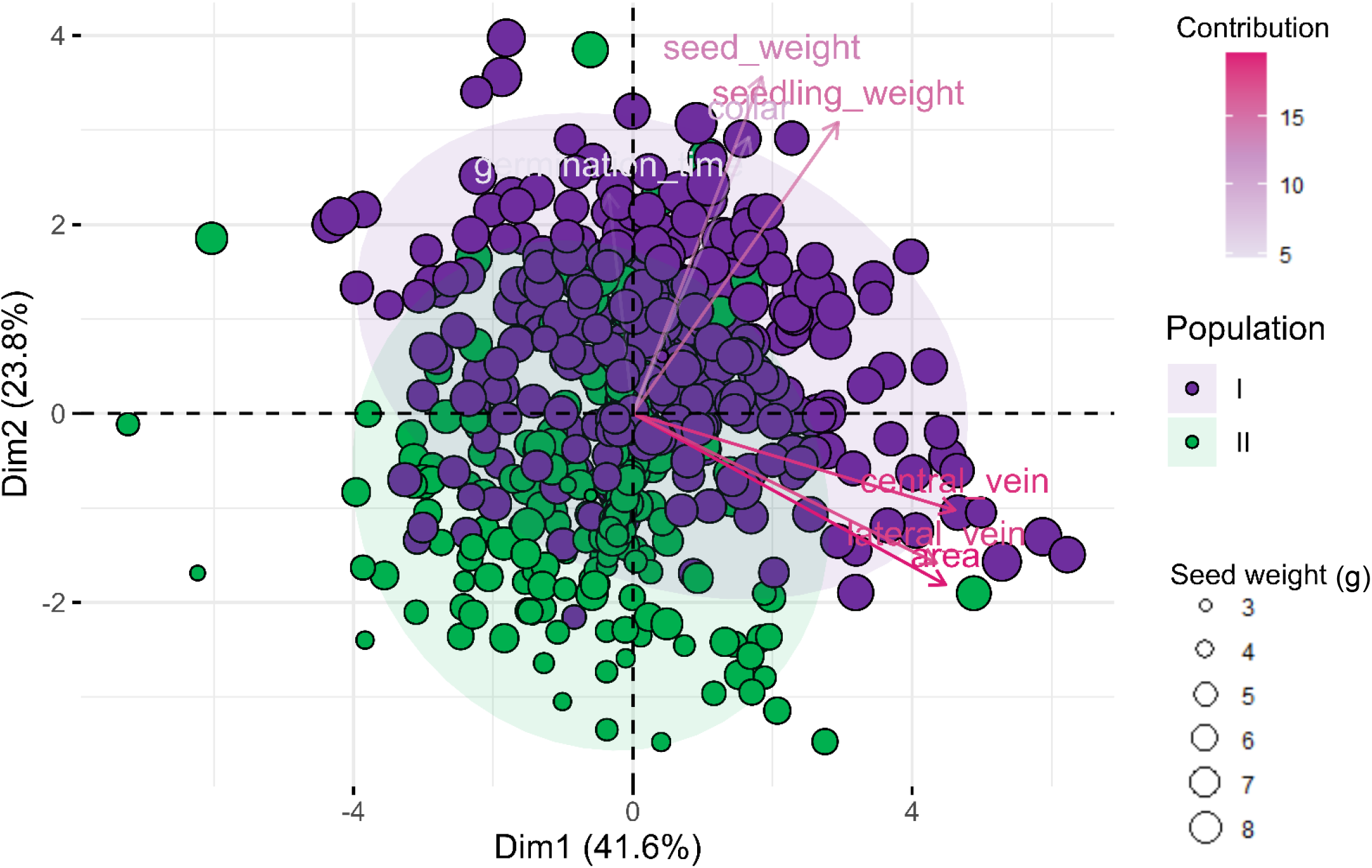
Principal component analysis (PCA) biplot of the seed and seedling measurements of *Hymenaea courbaril* according to the origin: Population I 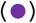 and Population II 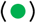. The size of the points is according to the seed mass. Vectors’ gradient colors are associated with their contribution to data variance. Concentration ellipses present a confidence interval of 95%.

The analyzed populations had different dispersion patterns (Figure 8). Population I had greater values for germination time, seed and seedling weight, having a greater confidence ellipse, and occupying a superior position in the principal component analysis (Figure 8). While the population II had consistent variations in seed weight and required less time to germinate.

Seed and seedling weight, and also collar diameter had similar directions (Figure 8) and occupied the same quadrant. This indicates that, in general, those features are more related to each other than with the protophilus dimensions, which were in the other quadrant, forming another group.

Biometric descriptions are essential to detect genetic variability between populations of a species and to identify morphologic traits to differentiate species from the same genus [4]. Principal component analysis (PCA) indicates how different populations are on the individual’s traits (Figure 8). The populations have differences in the position occupied by its individuals, and the studied measurements had different importance to describe these population traits. However, it is interesting to note that the concentration ellipses overlap each other at the center of the scatterplot, indicating that there is still a relative similarity between them.

In general, morphologic features presented by seedlings are related to their development and initial establishment in the natural environment [16]. The results also indicate that even when seeds are kept under the same environmental conditions, there are intrinsic differences in the responses of each individual, and when observed at the population scale, there is diversity between the groups (Figure 8). This could indirectly demonstrate the remaining genetic diversity of the populations, which could influence the usual wide divergence concering the best conditions for forest species to express their germination potential [16].

*H. courbaril* is an allogamous species, which means that this species favors cross-fertilization, tolerating only about 10% of self-fertilization [38]. This type of reproduction system results in great diversity inside the same population than when populations are compared [36], [38]. The PCA corroborates this hypothesis since there are great variations inside the same population, but when comparing the confidence ellipses, populations overlap each other (Figure 8). However, it is necessary to consider that some individuality remains between them.

The protophilus associated measurements were orthogonal to the other studied measurements (Figure 8), indicating a poor relationship between the collar diameter, seed, and seedling weight to the central and lateral vein length and protophilus area. As previously seen in Figure 6, the correlation between seed weight and protophilus dimensions and area was not strong.

### Protophilus elongation patterns

The protophilus of *H. courtoril* is described with a round apex and full edge. I16]. The leaf suffers a color change during the development [16], an event also observed in this study. This section aimed to understand how the greater protophilus areas were obtained.

Protophilus dimension development is not completely understood in this specific case. Hence, as the obtained data could provide some consistent information concerning the elongation pattern of the first leaf pair, and the possible relations between those variables (Figure 8).

The growing patterns proposed in this study (Figure 9) were based on the observed relations between the expansion of the protophilus area correlated with the increasing dimensions of the central vein (midvein) and lateral vein (leaf width). The proposed development patterns are presented in Figure 9.

**Figure 9:**
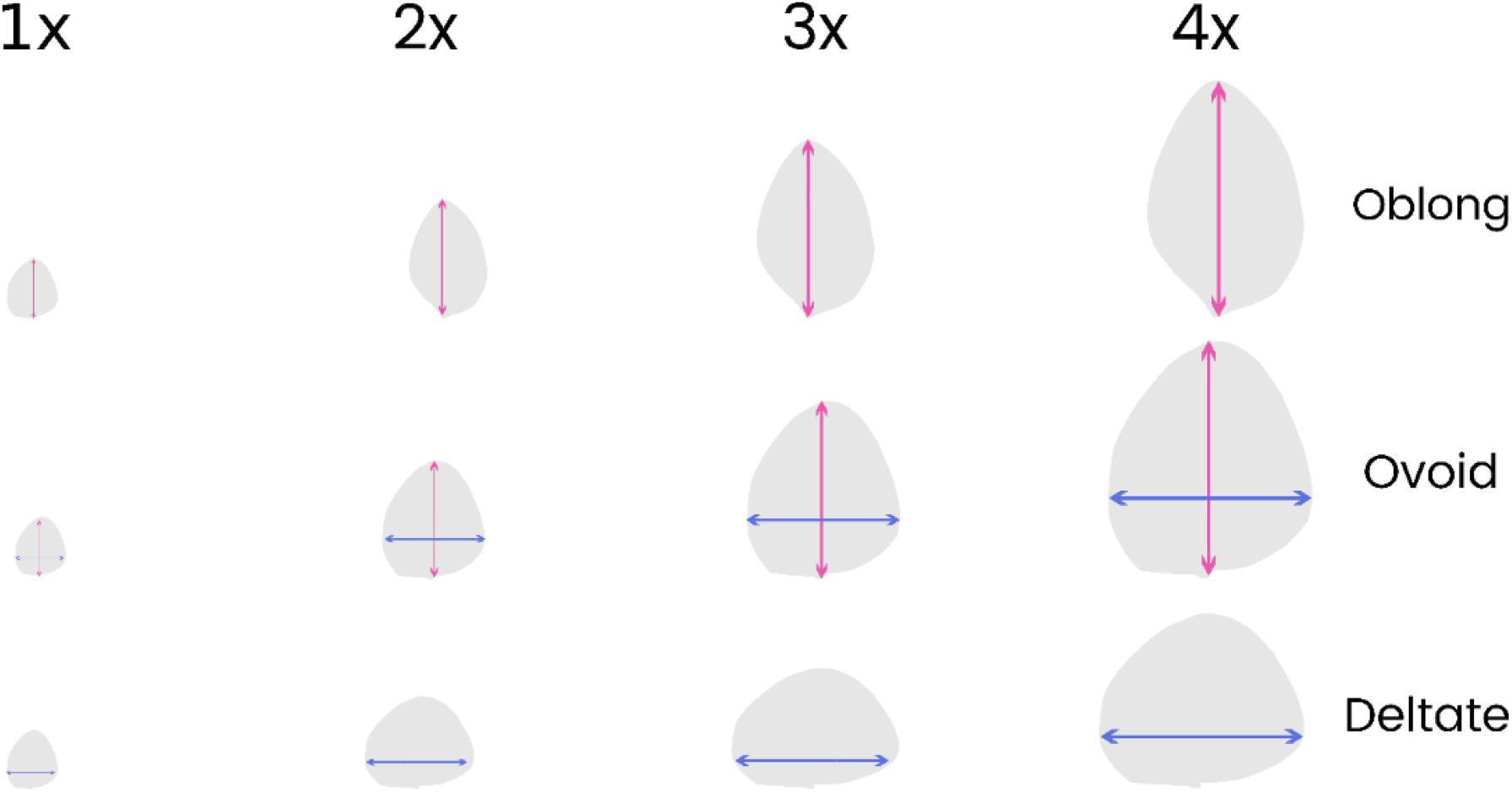
Protophilus theoretical elongation patterns: **Oblong** – elongation has mainly the central vein length 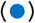 increase with leaf development; **Ovoid** – has central vein and lateral vein 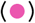 increasing at the same rate with elongation, and in **Deltate** – the protophilus have mainly lateral vein increasing with protophilus development.

The regression line that best describes the relation between the areas increases associated with the vein length is exponential (Figure 10), which is expected because the area is a squared measure and lengths are linear variables. Greater values of the protophilus area were obtained with the increase of lateral veins, while the central veins promoted a lesser increase in the area (Figure 10).

**Figure 10:**
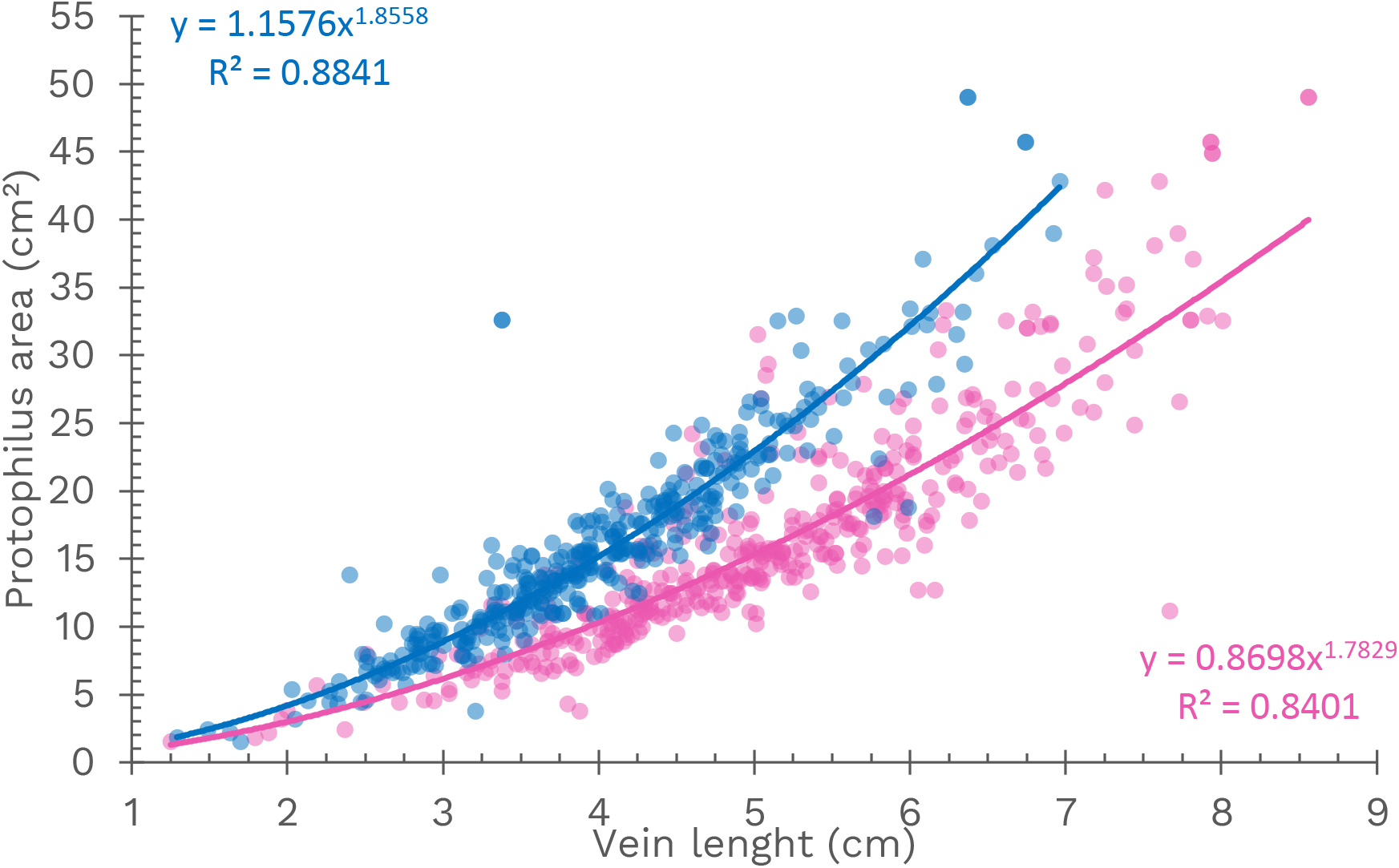
Scatter plot of the protophilus area according to the general regression line and its correlation indexes (R^2^) are also presented to the protophilus dimensions of *Hymenaea courbaril* Central vein 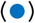 and Lateral vein 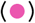.

The high correlation between the lateral vein and the central vein to the protophilus area could be observed previously in the principal component analysis (Figure 8). In the scatterplot, it is possible to understand how these measurements are associated with each other (Figure 10). Both regression lines were drawn using exponential regression, and the correlation index for both measurements is considered high [45], which is expected because the protophilus area is directly dependent on its dimensions.

The regression lines are very consistent with the smaller and intermediate values of the veins, but at the greater values, there is an increase in the uncertainty between the provided equation and the original data (Figure 10). Thus, the data indicate that a greater leaf area is not completely explained by the exponential model.

The growth of leaf blade is a complex process [50]. The leaf lamina starts its development early during the first stages of leaf formation, but expansion and differentiation occur during secondary morphogenesis [51]. The junction between the blade and the petiole region formation seems to be an essential step that determines the blade area at the blade-initiation stage [50]. Many factors could affect each step of the process evolved in leaf blade formation.

Recognizing the biometry of plants is very important for recovery programs because it makes recognition of the plants easier in the field and increases the understanding of normal development [4], [10]. The aim to using this sort of approach is to determine which of the studied dimensions is strongly associated with the protophilus area increase. Thus, it is clear that protophilus width has a considerable impact, when compared to midvein, with the increase of this leaf area, and presents more reliable regression values. Indicating that the real protophilus growing pattern is probably closer to the pattern that generates a deltate form (Figure 9). This association could be responsible for the similarities in the results mentioned in the Figure 5. Therefore, estimations of the leaf area that use the leaf width as a basic measurement would obtain more reliable values than using the central vein. However, caution is required because morphometric analysis traits consist of a continuum relation between the variables and not only a binary response [18].

It is also important to check how connected, both, the lateral and central veins, could be (Figure 11). The association between these variables is also strong but smaller than the values obtained for the correlation indices when those measures are associated with leaf area (Figure 10). Thus, the elongation of one dimension is also followed by the rise of the other, but not at the same taxa as the area, a square measure, is.

**Figure 11:**
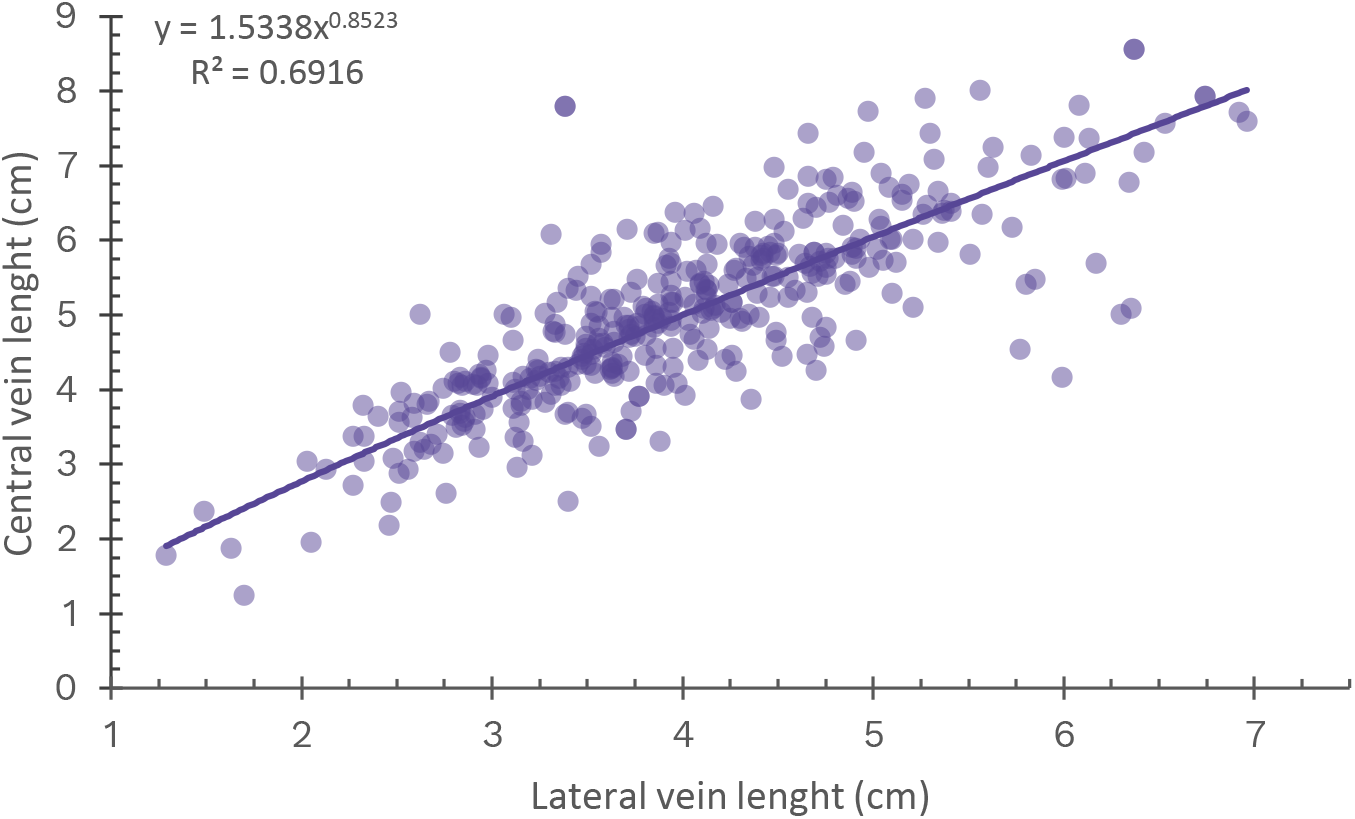
Scatter plot of protophilus lateral vein length and central vein length of *Hymenaea courbaril* A general regression line and its correlation indexes (R^2^) are also presented.

It is important to note that this data dispersion is most represented also through an exponential model, and not the linear, as expected (Figure 11). Indicating a not so simple or direct association.

The gene *an* in *Arabdopsis taliana* was responsible for reducing the leaf width, without affecting the leaf length [52], changing the proportions obtained for the wild type. In the first stages of leaf development, it seems that elongation starts in the leaf-length direction and then another gene group works together to promote leaf blade width growth [52]. Thereby, the development of protophilus measurements seems corroborate this hypothesis, in which there are different gene groups, but somehow associated with each other (Figure 11). It remains to be discovered how the activities of these different regulators of lamina initiation and growth are coordinated [53], and even if protophilus have a different development program for mature leaves.

## CONCLUSION

There was a consistent variation in protophilus’ central, vein length, seedling collar diameter, seed, and seedling weight according to population origin. Seed weight had a low association with the time required for germination, protophilus dimensions, and area. A medium correlation was found between seedling weight and collar diameter. The populations of *H. courbaril* have a relative similarity between them, but inside the same population, there were great discrepancies. Most of the individual variations were obtained to protophilus dimensions and area. The lateral vein was strongly associated with an increase in the protophilus area than the central vein.

## Acknowledgments

I would like to thank Victor Leite Jardim for his professional work and also Rafaela C. Perez, Juliana de Lemos, and Caio G. Tavares Rosa due to their consistent scientific support and encouragement.

